# Treatment with Metal-Organic Frameworks (MOFs) elicits secondary metabolite accumulation in *Aquilaria crassna* (Agarwood) callus culture

**DOI:** 10.1101/2024.08.23.609323

**Authors:** Sebastian Overmans, Yazan Alflayyeh, Sergio Gutiérrez, Yousef Aldlaigan, Kyle J. Lauersen

**Affiliations:** Bioengineering Program, Biological and Environmental Sciences and Engineering Division, King Abdullah University of Science and Technology (KAUST), Thuwal, Jeddah, Kingdom of Saudi Arabia; Plant Tissue Culture & Biotechnology Center, Ministry of Environment Water and Agriculture of Saudi Arabia (MEWA). P.O.Box 11195, Riyadh, Kingdom of Saudi Arabia

**Keywords:** Agarwood, *Aquilaria crassna*, Metal-Organic Frameworks (MOFs), plant callus, tissue culture, secondary metabolites, sustainability

## Abstract

Thymelaceaous trees are prized for accumulating fragrant resins composed of hundreds of secondary metabolites in their woody tissues. Slow growth and increasing consumer demand have stretched natural sources of agarwood trees to being endangered and alternative production modes, including silviculture and tissue culture, are currently being investigated. Dedifferentiated tissue culture of agarwood trees provides a means of cell propagation independent of environmental context. However, secondary metabolite accumulation, as found in fragrant resins, occurs largely in response to wounding. Here, we investigated the application of metal-organic frameworks (MOFs) as potential elicitors of secondary metabolite formation in *Aquilaria crassna* tissue culture samples. Callus cultures were exposed to five commercially available MOFs: UiO-67, MOF-808, HKUST-1, ZIF-67, and MOF-74, and ethanol extracts were used to quantify secondary metabolite accumulation compared to untreated cultures. Samples that were exposed to Zr-based MOFs exhibited similar metabolite production profiles, (trans-2-Carboxy-cyclo-hexyl)-acetic acid was reduced in the presence of all MOFs, the Cu-containing HKUST-1 MOF increased palmitic acid levels, and MOF-808 and ZIF-67 were found to elicit the highest accumulation of secondary metabolites with potential fragrance applications. These results demonstrate the possibility of eliciting secondary metabolites from dedifferentiated agarwood tree cell culture and may provide an alternative means of sourcing fragrant specialty chemicals from these plants.

## 1 Introduction

*Aquilaria* sp. and other trees belonging to the family Thymelaeaceae are of high economic and cultural value due to the aromatic resin produced within their heartwoods after wounding (Hishamuddin *et al*., 2019). Their resinous fragrant wood, ‘agarwood’, is extensively used in fragrances, traditional medicine, and as incense across various cultures (López-Sampson and Page, 2018; Shivanand *et al*., 2022). The formation of agarwood involves accumulating a variety of secondary metabolites produced in response to stressors, such as physical injury or microbial infection (Ma *et al*., 2023; Shivanand *et al*., 2022). These metabolites comprise a broad spectrum of chemicals, including sesquiterpenoids, 2-(2-phenylethyl)chromones (PECs), alkaloids, flavonoids, phenolic acids, and triterpenoids among others. These compounds collectively create the unique aromatic and therapeutic properties of agarwood (Gao *et al*., 2019; Gutiérrez *et al*., 2024b; He *et al*., 2022).

Global demand for agarwood resin has been increasing, with first-grade samples costing as much as US$ 100,000 per kilogram (Lee and Mohamed, 2016), while natural reserves are strained due to the slow growth of the trees and inconsistent localized resin accumulation around points of wounding. Recent reports show that demand for agarwood has outpaced the supply of natural resources to the point that the Convention on International Trade in Endangered Species of Wild Fauna and Flora (CITES) has declared *Aquilaria* spp. as potentially threatened with extinction (CITES, 2024). Exploration of alternative and more sustainable methods to produce *Aquilaria* secondary metabolites is needed (Gutiérrez *et al*., 2024b; Shivanand *et al*., 2022). Various methodologies have been deployed to artificially induce controlled resin formation in agarwood trees (Tan *et al*., 2019). One approach involves using callus cultures derived from *Aquilaria* sp. shoot segments. When subjected to abiotic stress, these cultures have been observed to exhibit an enhanced production of aromatic terpenoids (Zhang *et al*., 2021). Various techniques, such as the application of methyl jasmonate, mechanical wounding, and fungal inoculation, have been tested for their potential to induce the requisite stress in plant tissues, with variable outcomes (Ngadiran *et al*., 2023; Tan *et al*., 2019; Zang *et al*., 2016).

An approach that has gained attention in recent years is the use of Metal-Organic Frameworks (MOFs). MOFs are crystalline materials composed of metal ions or clusters interconnected by organic ligands that form porous structures and have many applications (James, 2003), such as controlled chemical release (Chauhan *et al*., 2022; Rojas *et al*., 2022; Wang *et al*., 2023). MOFs have large surface areas, high porosities, and tunable chemical properties, and are used in diverse fields, including agriculture, gas storage, catalysis, and drug delivery (Abdelhameed *et al*., 2019; Niknam *et al*., 2018; Rojas *et al*., 2022; Zhou *et al*., 2012). In living organisms, MOFs can act as nanoparticles that physically interact with cells and consequently can elicit stress responses (Al-Rehili *et al*., 2019; Chauhan *et al*., 2022; Guan *et al*., 2021; Hu *et al*., 2024; Rojas *et al*., 2022).

Here, we aim to evaluate the ability of MOFs to induce abiotic stress in *Aquilaria crassna* callus samples, and determine their efficacy in eliciting secondary metabolite accumulation typically observed following physical wounding of mature tree tissue. Five different commercially available MOFs were tested at various concentrations, and changes in the metabolite profiles of *A. crassna* callus samples following MOF application were analyzed. Overall, we observed specific differences in the secondary metabolite production dependent on the type and concentration of MOF. The results presented here highlight the high potential of metal-organic frameworks to potentially offer a new avenue to source agarwood secondary metabolites sustainably from cell cultures rather than mature trees.

## 2 Materials and Methods

### 2.1 Media and plant growth regulators (PGRs) for callus induction and proliferation

The initial *Aquilaria crassna* plant material was provided by the Ministry of Environment, Water, and Agriculture (MEWA) in Saudi Arabia as *in vitro* cultures. Callus cultures were induced from shoot segments, using a modified MS medium based on the formulation by Murashige and Skoog (Murashige and Skoog, 1962). To initiate callus formation and proliferation, the medium was enriched with 3% sucrose, 0.6% agar, 2.2 µM of 2,4-dichloro phenoxy acetic acid (Sigma-Aldrich, St. Louis, USA), and 2.3 µM of 6-benzyl amino purine (Sigma-Aldrich, St. Louis, USA) (Fig. 1). These specific plant growth regulators (PGRs) are known to be pivotal for plant cell differentiation and growth (Di Mambro *et al*., 2017; Gaba, 2005; Sabagh *et al*., 2021), and have been used for callus induction and proliferation in various plant species, including those belonging to the genus *Aquilaria* (Jayaraman *et al*., 2014; Qinying *et al*., 2001). To provide a suitable environment for growth and development, all callus cultures were incubated at 25 ± 1 ºC in darkness for 8 weeks before being used for subsequent experiments.

**Figure 1.**
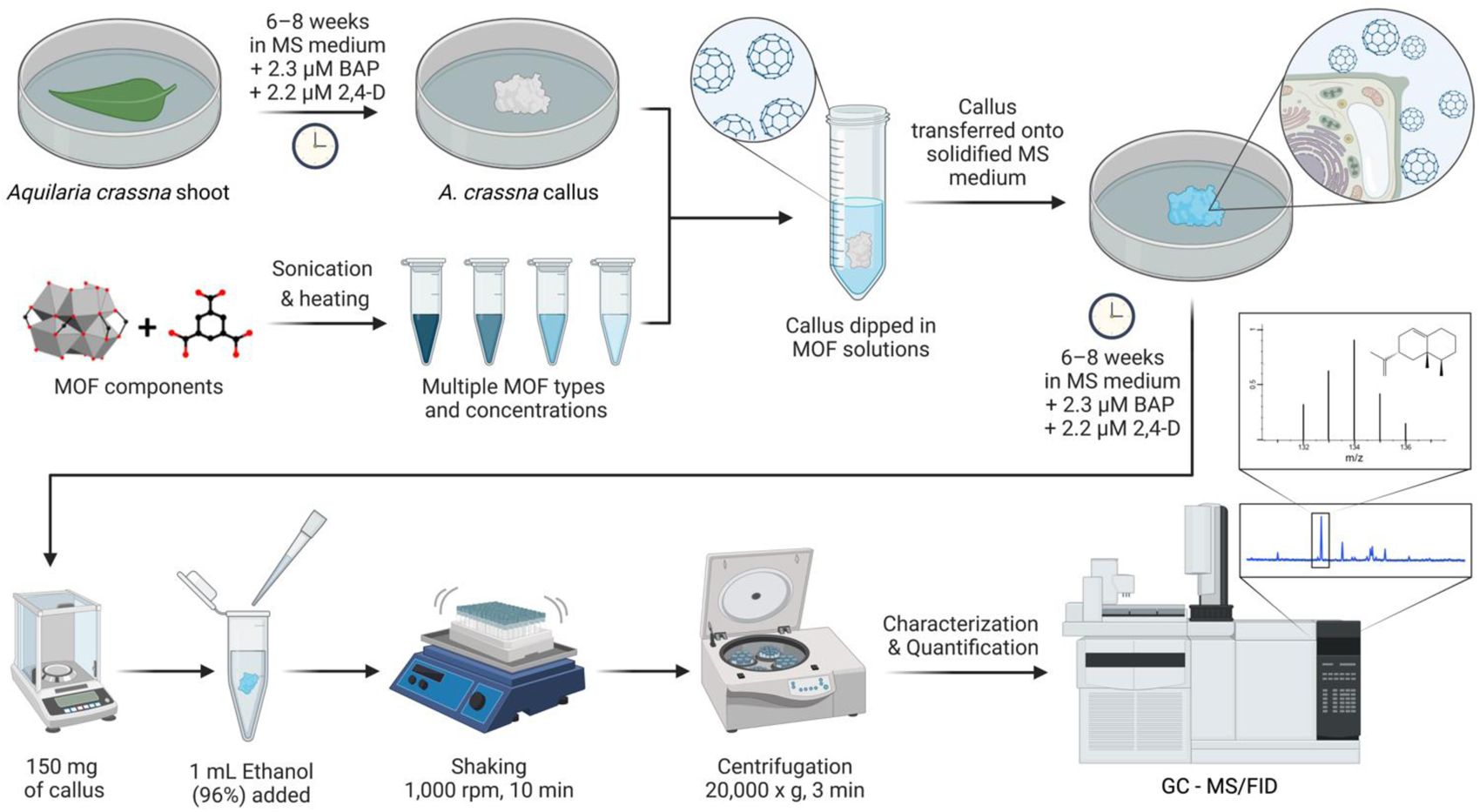
Workflow of callus cultivation, MOF treatment, and subsequent sample processing and specialty metabolite GC-MS/FID quantification/identification that were used in the present study. Figure was created with BioRender.com.

### 2.2 Determination of suitable solvent for *Aquilaria* metabolite extraction

Before extracting the metabolites from MOF-treated callus samples, a preliminary experiment was conducted to identify the most suitable solvent, characterized by a high extraction efficiency and low GC-MS background signal. Approximately 500 mg of dried and ground *Aquilaria* wood sample was weighed and transferred into each of nine individual microcentrifuge tubes to which 1 mL of one of the following nine solvents was added: methanol, 96% ethanol, acetone, dichloromethane (all from VWR International, Fontenay-sous-Bois, France), n-hexane (Acros Organics, Geel, Belgium), n-dodecane, tetrahydrofuran (both from Sigma-Aldrich, St. Louis, USA), and the two perfluorocarbons FC-770 (Fluorochem, Glossop, UK) and FC-3283 (Acros Organics, Geel, Belgium). The mixtures of wood samples and solvents were briefly vortexed and then shaken at 200 rpm for 2 h at room temperature to facilitate the extraction of metabolites. Afterward, the samples were centrifuged at 8000 x *g* for 15 min to separate all solids from the liquid phase. 150 µL of each supernatant was transferred into a separate amber GC glass vial. The supernatants and blanks of each extraction solvent were analyzed on a gas chromatograph as described below.

### 2.3 Abiotic stress induction in callus cultures using MOFs

To simulate the natural wounding in Aquilaria trees that triggers the production of specialty metabolites, we induced abiotic stress in the callus cultures through exposure to five distinct MOFs (UiO-67, MOF-808, HKUST-1, ZIF-67, MOF-74) that were synthesized at and provided by King Fahd University of Petroleum and Minerals (Dhahran, Saudi Arabia) (Table 1). The selection of these MOFs was based on their commercial availability, varying pore sizes, stability, and chemical functionalities which we postulated could influence the extent of abiotic stress induced in the callus cultures. To address the low solubility of MOFs, firstly slurries were prepared before suspensions of each MOF were prepared at varying concentrations in distilled water (10–40 g L^−1^) (see Table 1), and autoclaved before further use (Fig. 1).

**Table 1.** Codes and properties of MOFs used in the study. Displayed are the MOFs’ identification codes, molecular formulas, CAS numbers, metals, ligands, colors, and the concentrations of MOF solutions tested in the present study.

| MOF ID | Molecular Formula | MOF CAS No. | Metal | Ligand | Color/code | Concentration (g/L) |
| --- | --- | --- | --- | --- | --- | --- |
| UiO-67 | C<sub>84</sub>H<sub>52</sub>O<sub>32</sub>Zr<sub>6</sub> | 1072413-83-2 | Zr | 4,4'-Biphenyldicarboxylic acid (CAS: 787-70-2) | white/ W1 | 10, 20 |
| MOF-808 | C<sub>24</sub>H<sub>16</sub>O<sub>32</sub>Zr<sub>6</sub> | 1579984-19-2 | Zr | Trimesic acid (CAS: 554-95-0) | white/ W2 | 10, 20, 30 |
| HKUST-1 | C<sub>18</sub>H<sub>12</sub>Cu<sub>3</sub>O<sub>15</sub> | 222404-02-6 | Cu | Trimesic acid (CAS: 554-95-0) | blue/ B | 10, 20, 30, 40 |
| ZIF-67 | C<sub>8</sub>H<sub>12</sub>N<sub>4</sub>.Co | 46201-07-4 | Co | 2-Methylimidazole (CAS: 693-98-1) | purple/ P | 10, 20 |
| MOF-74 | C<sub>8</sub>H<sub>4</sub>O<sub>8</sub>Mn<sub>2</sub> | 1235342-69-4 | Mn | 2,5-Dihydroxyterephthalic acid (CAS: 610-92-4) | red/ R | 10, 20 |

Four replicates of 8-week-old callus samples (n=4) were immersed in the MOF slurry, maintaining homogeneity through continuous shaking. All tools and containers were sterilized, and the procedure was conducted under aseptic conditions in a laminar flow hood. After immersion, the cultures were transferred onto solidified MS medium in Petri dishes and incubated in darkness at 25 ± 1 °C for an additional 8 weeks to facilitate their growth and development (Fig. 1).

### 2.4 Callus recovery and processing for metabolite analysis

After 8 weeks of cultivation, the petri dishes were photographed (Fig. 2), and the callus samples were recovered to assess the impacts the MOFs had on their growth and metabolite production. From each of the four replicate calluses per treatment (n=4), one sample of approximately 150 mg wet weight was cut, weighed, and immersed in 1 mL of 96% ethanol. To ensure an efficient extraction of metabolites, the mixtures were agitated at 200 rpm for 8 h. Subsequently, the suspensions were centrifuged at 8000 x g for 15 min to segregate the liquid phases from the solid residues. 150 µL of each supernatant was pipetted into amber glass vials for further GC-MS analysis.

**Figure 2.**
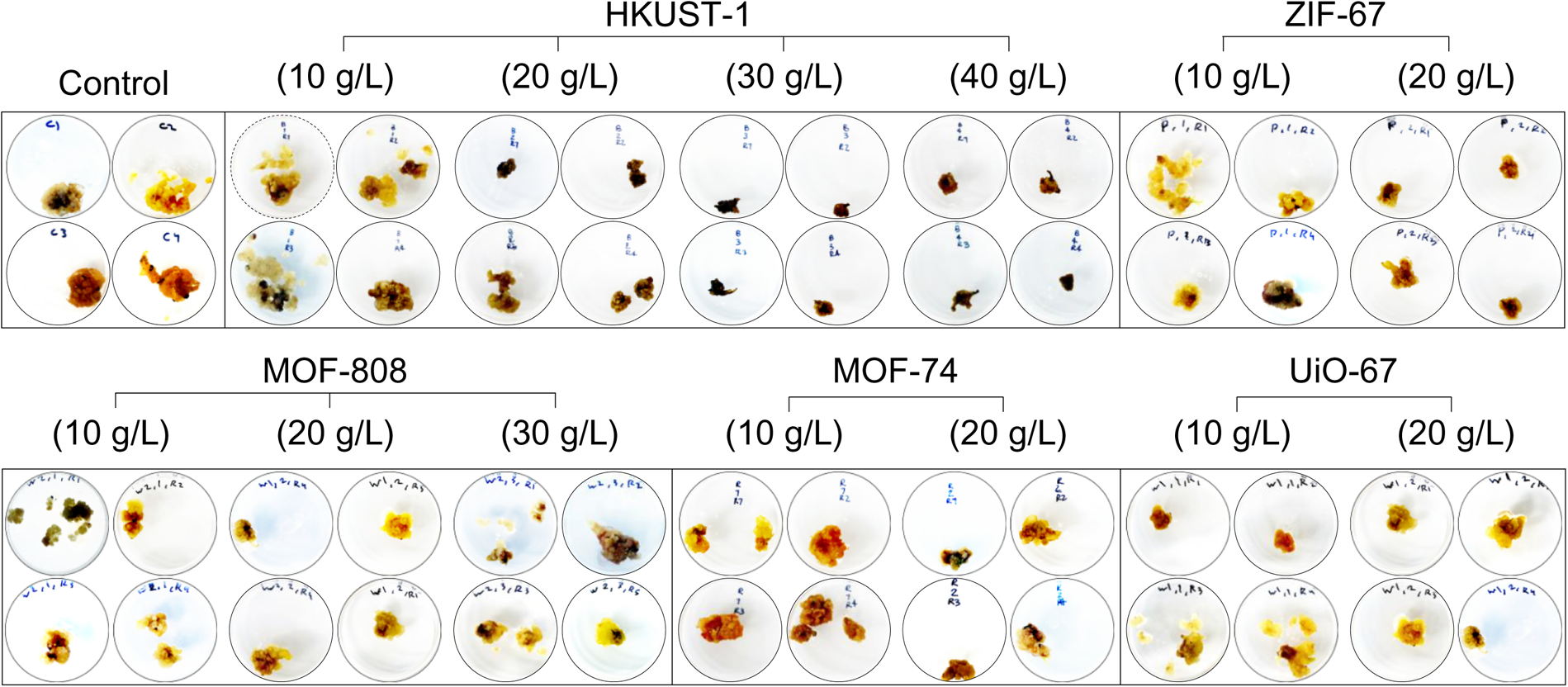
Photographs of callus samples that were briefly dipped in one of five different types of MOFs at varying concentrations (10–40 g/L), and of callus samples that were not exposed to any MOF (Control). All photographs were taken after 8-week cultivation on solidified MS growth medium. For each treatment, four replicate callus samples are shown (n=4).

### 2.5 Gas chromatography-mass spectrometry (GC-MS) analysis

All solvent samples were analyzed using an Agilent 7890A gas chromatograph (GC) coupled with a 5975C mass spectrometer (MS) with a triple-axis detector (Agilent Technologies, USA). The GC was equipped with a DB-5MS column (Agilent J&W, USA), with helium as the carrier gas at a flow rate of 1 mL per min. A previously described GC oven temperature protocol was used (Overmans and Lauersen, 2022). The analysis was conducted using a splitless injection to ensure maximum sensitivity. After a 13-min solvent delay, mass spectra were recorded across a scanning range of 50–750 m/z at a rate of 20 scans per second.

Chromatograms were processed and integrated using the MassHunter Workstation software v. B.08.00 (Agilent Technologies, USA). Metabolites were identified by comparing mass spectra against the National Institute of Standards and Technology (NIST) library (Gaithersburg, MD, USA), and standard mixtures of terpenoids were used as internal quality controls for the analysis. The peak areas of metabolites were normalized to the respective sample weight to account for variations in sample size. All GC-MS measurements were performed in technical duplicates (n=2), with manual verification of chromatograms conducted for quality control.

### 2.6 Data analysis

All data analyses and visualizations were performed using JMP version 16 (SAS Institute Inc, NC, USA) and GraphPad Prism v. 10 (GraphPad Software, MA, USA). Callus photographs were processed and white-balance corrected using Affinity Photo v. 1.10.6 (Serif Ltd., West Bridgford, UK). Visual elements were organized and harmonized using Affinity Publisher v. 1.10.6 (Serif Ltd., West Bridgford, UK).

## 3 Results & Discussion

### 3.1 Optimum solvent for extraction of *Aquilaria* metabolites

Before extracting metabolites from the MOF-treated callus samples, a preliminary experiment using dried agarwood was conducted to ascertain the most efficient extraction solvent. The optimum solvent was characterized by high extraction efficiency and minimal GC signal interference at the retention times of the target compounds. Within the retention time range of 14–26 min, where the compounds of interest were expected, 5 solvents: dichloromethane (DCM), dodecane, FC-770, hexane, and tetrahydrofuran (THF) exhibited many background peaks (Fig. 3), rendering them unsuitable for subsequent metabolite extraction due to the complication in identifying target compounds. Among the remaining 4 solvents, FC-3283 demonstrated inefficient metabolite extraction from agarwood, as evidenced by the low diversity and amounts of compounds in the solvent extracts. The relatively lower extraction efficiency of FC-3283 compared to traditional alkanes such as dodecane has recently been discussed (Gutiérrez *et al*., 2024a; Overmans and Lauersen, 2022). Comparative analysis of chromatograms from acetone, ethanol, and methanol extracts revealed similar profiles between those solvents. Overall, ethanol was identified as the most suitable solvent for further experiments based on its superior efficiency in extracting a diverse and abundant range of agarwood-derived metabolites, as observed from the chromatograms (Fig. 3). This finding is in line with a recent microalgae study in which ethanol was also used as the final solvent to capture terpenoid metabolites typically produced by *Aquilaria* sp. (Gutiérrez *et al*., 2024b). Acetone extracted approximately the same quantities of metabolites as ethanol but was not used for further experiments because it is considered more hazardous and corrosive than ethanol (de Jesus and Filho, 2020; Zou *et al*., 2021).

**Figure 3.**
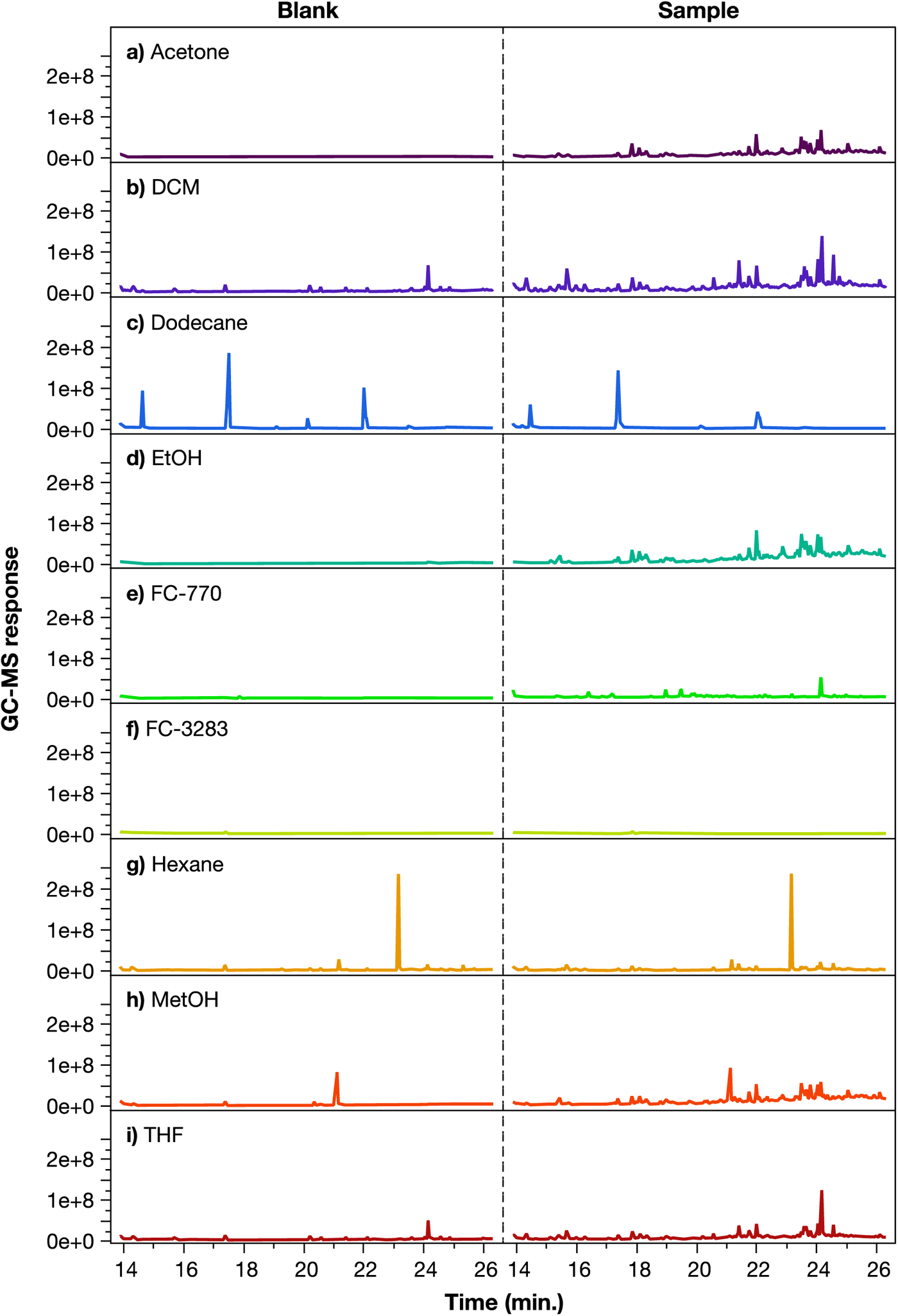
GC chromatograms of pure solvents (left panels) and of agarwood extracts (right panels) from the preliminary extraction-solvent experiment with acetone, dichloromethane (DCM), dodecane, ethanol (EtOH), FC-770, FC-3283, hexane, methanol (MetOH), and tetrahydrofuran (THF).

### 3.2 Effect of MOF exposure on secondary metabolite production in *Aquilaria* callus cultures

Exploring the use of metal-organic frameworks (MOFs) to enhance secondary metabolite production in *Aquilaria* species revealed varied effects based on the specific MOF type used. Overall, the MOFs UiO-67 and UiO-74 were found to be less effective in inducing the production of secondary metabolites (Fig. 4). In contrast, exposure to MOFs 808 and ZIF-67 resulted in the highest production of secondary metabolites (Fig. 4). These findings are in agreement with previous studies demonstrating that exogenous metals can significantly alter plant metabolite profiles under stress conditions (Liu *et al*., 2024; Parwez *et al*., 2023; Yang *et al*., 2024). Secondary metabolites with applications in consumer fragrance products such as aristoline, geranyl isovalerate, and juniperol were not at all present in the experimental controls although were detected in some MOF-treated samples (Fig. 5a). The same was observed for the lactone 4-octylbutan-4-olide and pipradrol, which were both only produced by the callus samples following treatment with any MOF. Pipradrol and its derivatives have been studied intensively for their pharmacological potential (Liechti *et al*., 2014), as they are proposed to enhance neurotransmitters like dopamine and norepinephrine in the brain (White and Archer, 2013).

**Figure 4.**
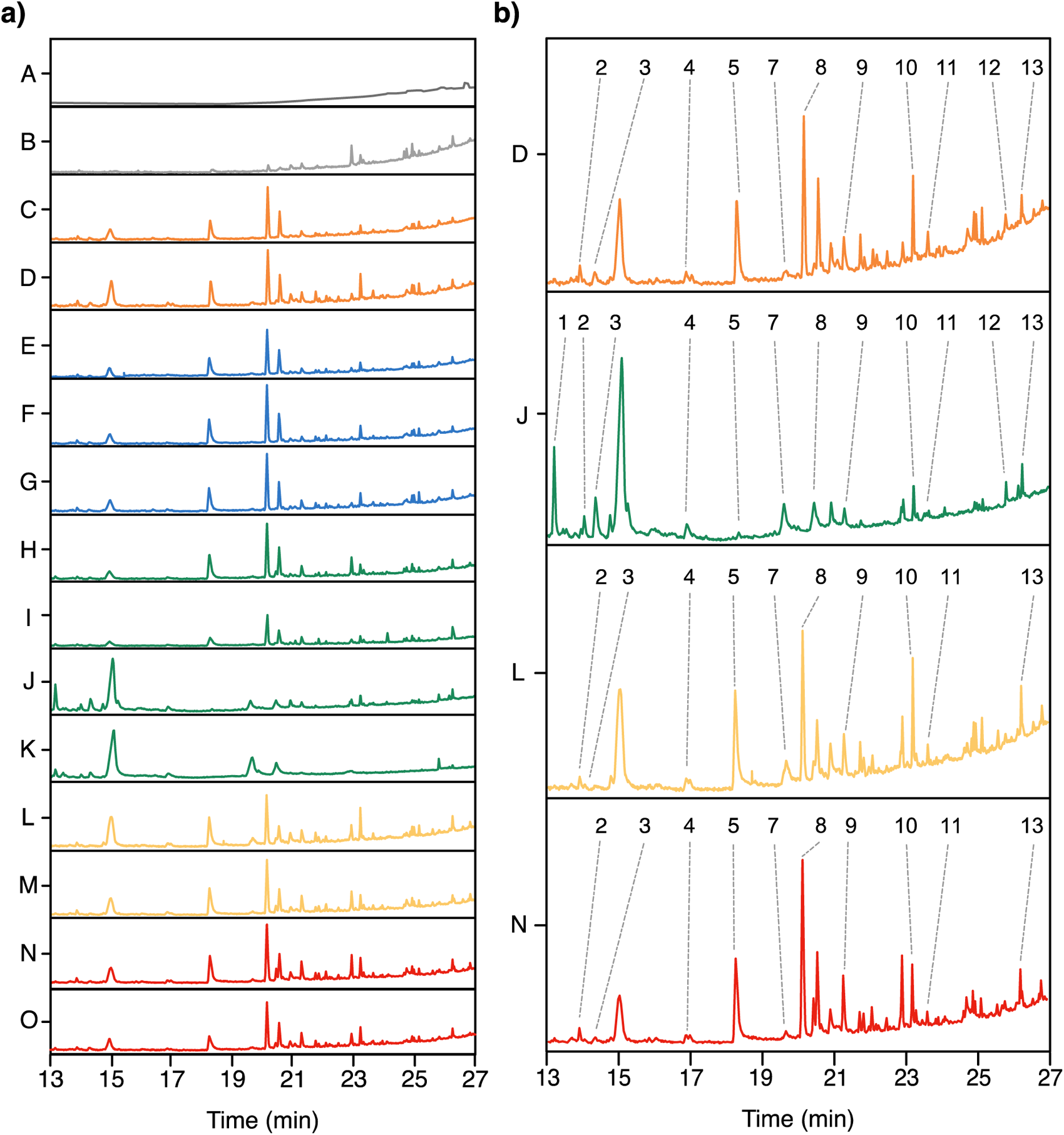
GC-MS chromatograms and metabolite identification. **a)** Representative GC-MS chromatograms showing retention times of metabolites from *A. crassna* callus tissue exposed to different MOFs. Chromatogram annotation code: **(A)** Ethanol blank; **(B)** Negative control (no MOF); **(C, D)** UiO-67 treatments (10–20 g/L); **(E, F, G)** MOF-808 treatments (10–30 g/L); **(H, I, J, K)** HKUST-1 treatments (10–40 g/L); **(L, M)** MOF-74 treatments (10–20 g/L); **(N, O)** ZIF-67 treatments (10–20 g/L). **b)** Selection of chromatograms highlighting peaks of metabolites identified after comparison against the NIST Library. Identified metabolites and retention times (RT) include: **(1)** Pipradrol (RT 13.2), **(2)** Spathulenol (RT 14.3), **(3)** Aristoline (RT 14.6), **(4)** Butylated hydroxyanisole (RT 16.8), **(5)** trans-2-Carboxy-cyclohexyl-acetic acid (RT 17.9), **(6)** Geranyl isovalerate (RT 18.2), **(7)** Longicamphor (RT 19.5), **(8)** 4-Octylbutan-4-Olide (RT 20.4), **(9)** Valencene (RT 21.2), **(10)** Juniperol (RT 24.3), **(11)** Longipinane (RT 24.6), **(12)** Hexadecanoic acid, ethyl ester (RT 26.3), **(13)** Palmitic acid (RT 26.8).

**Figure 5.**
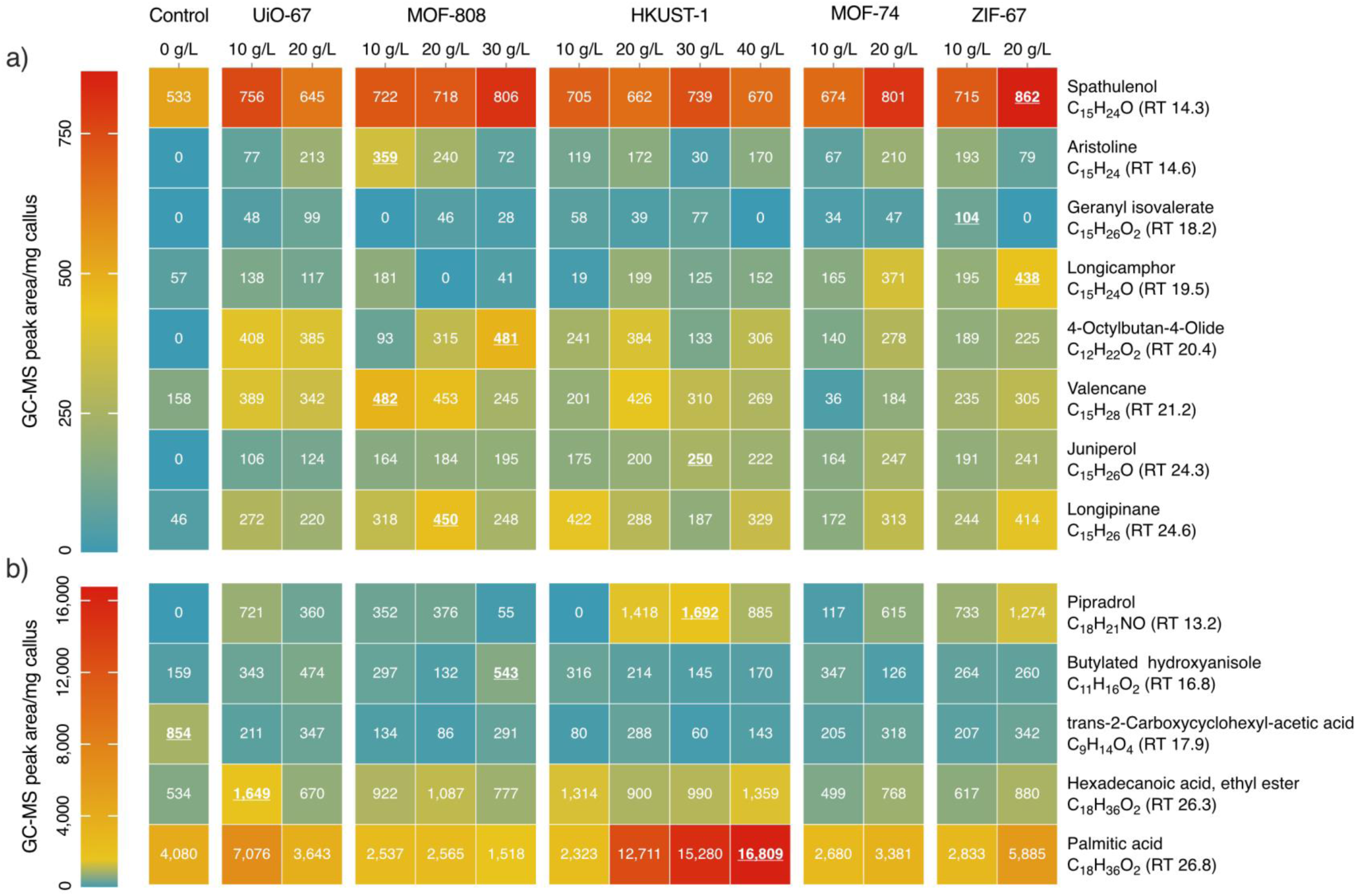
Relative quantification (as GC-MS peak area/mg callus) of various **a)** fragrant metabolites and **b)** other prominent compounds produced by *A. crassna* callus cultures following different MOF treatments. RT refers to the GC retention time in minutes. Numbers inside boxes represent mean values of four replicates (n=4). The highest value in each row is underlined.

Other high-value compounds, including the terpenoids spathulenol, longicamphor, valencane, and longipinane were present at low concentrations in untreated callus samples. However, exposure to MOFs resulted in concentration increases of up to 10-fold as in the case of longipinane when callus was treated with MOF-808 (Fig 5a). The presence of any MOF led to a reduction in trans-2-carboxycyclohexyl-acetic acid (Fig. 5b). While its potential application remains unclear, this compound has been identified in various studies, notably in the context of natural products and their biological activities. For instance, this chemical has been recognized as a significant component of fractions derived from the bark of the tropical tree *Alstonia boonei*, which is has been investigated for its anti-obesity and anti-lipolytic effects (Anyanwu *et al*., 2018).

The MOF HKUST-1, which contains copper as the metal base, greatly enhanced the production of palmitic acid by up to 4-fold compared to controls (Fig. 5b). Palmitic acid is a major component of agarwood (Aqmarina Nasution *et al*., 2020; Ogita *et al*., 2015; Wang *et al*., 2018). An elevated production of this compound indicates the onset of a defense mechanisms involving increased formation of free fatty acids that can trigger oxidative burst and fatty acid oxidation cascades in response to an external stressors (Sen *et al*., 2017).

Samples exposed to MOFs UiO-67 and MOF-808 which contain zirconium exhibited similar metabolite production profiles (Fig. 4; Fig. 5). This metal has no essential functions in plant metabolism and is generally considered to be of low toxicity (Shahid *et al*., 2013). However, it has been reported that zirconium can reduce the growth of wheat plants and affect their enzyme activity (Fodor *et al*., 2005), which may explain why callus samples treated with MOFs containing this metal exhibited a perturbed metabolite profile here.

We observed no clear trend across MOFs or compounds with regards to the effect of MOF concentration on secondary metabolite production. However, secondary metabolite accumulations were strongly dependent on MOF type and their applied concentration. For example, palmitic acid and valencene decreased with increasing concentrations of UiO-67 and MOF-808 (Zr-based), whereas with MOF-74 (Mn-based) and ZIF-67 (Co-based), more valencene was produced at higher MOF concentrations (Fig. 5a). However, this pattern of gradual changes in production with increasing concentration is not consistent across all compounds. For some secondary metabolites, their production was highest at intermediate MOF concentrations. This finding indicates MOF type and concentration need to be carefully selected to maximize the production of a given target compound without leading to callus senescence.

### 3.3 Conclusion

The use of MOFs for the production of secondary metabolites from the callus of *Aquilaria* sp. presents a promising avenue for future research. Systematic empirical testing of MOFs with *Aquilaria* sp callus and its subsequent metabolite analysis can enable stress-elicitation to obtain valuable compounds from this plant. However, whether this process could be effectively scaled to produce consumer products, like fragrances, remains to be seen. This approach could help improve the sustainability the consumer fragrance industry but also holds potential for applications in pharmaceuticals, agriculture, and other industries requiring plant-derived compounds.

## Author contributions

Y ALF performed the experiments, contributed to the research planning, experimental design, data collection and visualization, formal analysis, and reviewed the manuscript. SO and SG contributed to the research planning, experimental design, GC-MS analysis, and data visualization, and wrote the original manuscript draft. The research was conducted in the laboratories of Y ALD and KJL, who were responsible for experimental design, project scope, funding acquisition, supervision, and revision of the manuscript. All authors have read and approved the final manuscript version.

## Acknowledgments

The authors would like to thank Sultan Al Gethami & Dr. Ko Chien Ying at the Plant Tissue Culture & Biotechnology Center for their assistance with the cultivation of plant material, and Islam Tayeb at KFUPM for synthesizing the MOFs used in this study. The research reported in this publication was supported by KAUST baseline funding awarded to Kyle Lauersen, and through financial support from the Saudi Ministry of Environment Water & Agriculture (MEWA). Yazan Alflayyeh’s research stay at the Saudi Ministry of Environment Water & Agriculture (MEWA) was organized by King Abdulaziz and His Companions Foundation for Giftedness and Creativity (Mawhiba).

